# Attention selectively gates afferent signal transmission to area V4

**DOI:** 10.1101/019547

**Authors:** Iris Grothe, David Rotermund, Simon David Neitzel, Sunita Mandon, Udo Alexander Ernst, Andreas Kurt Kreiter, Klaus Richard Pawelzik

## Abstract

Selective attention causes visual cortical neurons to act as if only one of multiple stimuli are within their receptive fields. This suggests that attention employs a, yet unknown, neuronal gating mechanism for transmitting only the information that is relevant for the current behavioral context. We introduce an experimental paradigm to causally investigate this putative gating and the mechanism underlying selective attention by determining the signal availability of two time-varying stimuli in local field potentials of V4 neurons. We find transmission of the low frequency (<20Hz) components only from the attended visual input signal and that the higher frequencies from both stimuli are attenuated. A minimal model implementing routing by synchrony replicates the attentional gating effect and explains the spectral transfer characteristics of the signals. It supports the proposal that selective gamma-band synchrony subserves signal routing in cortex and further substantiates our experimental finding that attention selectively gates signals already at the level of afferent synaptic input.

## Introduction

The sense organs deliver a continuous stream of comprehensive information about the environment. Selective attention is required to focus on parts of the incoming information, thereby allowing for advanced processing like shape perception of individual stimuli embedded in cluttered visual scenes (1–3). In a now classical experiment aiming to demonstrate such selective processing on the neuronal level, Moran and Desimone (4) placed either one or two stimuli in a visual neuron’ s receptive field (RF). They found for neurons in visual areas V4 and IT, that when a preferred and a non-preferred stimulus were placed together in the RF, the firing rate response of the neurons was intermediate compared to the responses when the stimuli were presented alone. However, when attention was allocated to just one of the two stimuli, the neurons responded as if only this attended stimulus was present in its RF. This result and succeeding studies (5–8) all using simple, mostly static stimuli with opposed properties in feature space, provide evidence suggesting that attention can gate visual signals. Still, so far nobody has been able to show signal gating explicitly. We introduce a method, which does not rely on feature differences between simple stimuli, to prove attentional signal gating in a rigorous sense: for arbitrary, time-varying signals. Further, the neuronal mechanisms underlying such signal gating are controversial. Two major classes of proposed mechanisms can be distinguished, which either selectively gate transmission of neuronal signals already at the level of afferent inputs or allow in contrast all afferent signals to enter local processing in a similar way and attention rather modifies the local computations by modulating the neurons output. In the first class of proposed mechanisms neurons respond “as if” only the attended stimulus was present because the corresponding group of afferent inputs from upstream areas is selectively processed. This might be achieved by differential modulation of the synaptic input, i.e. selectively changing the efficacy of signal transmission for subsets of afferent inputs, without necessarily changing spike rate in the afferent population from which the afferent inputs derive. As a result of such selective input gating, local neuronal processing would be predominately determined by signals representing the attended stimulus (e.g. 8, 9–11). The second class assumes that attention could act directly on the response strength of the neurons. In contrast to the first, this explanation suggests that all signals representing the two stimuli enter into local neuronal processing independent of the direction of attention. Here, the neuronal response “as if” only the attended stimulus is present results from a differential modulation of the output gain of neurons distinguished by their stimulus selectivity (12–17). We set out to directly assess whether attention is capable of switching neuronal processing between input signals as assumed by differential input gain modulation or whether the observed rate effects result rather from output gain modulation. Distinguishing these two mechanisms is inherently difficult since the spiking rate output is reflecting only the final outcome of the neuronal computations with unknown contributions from the different input signals. Therefore, we constructed an experimental paradigm that allows to infer simultaneously and independently to which extent two stimuli placed within the same receptive field are causally influencing neuronal activity recorded in visual cortex. We found that the contribution of the attended stimulus to local neuronal processing was strongly enhanced, showing that arbitrary, time-varying signals can be selectively gated by attention and suggesting a strong selection of specific inputs for selective processing. Furthermore, simulations demonstrated that such input signal gating could be mechanistically implemented by gamma-band synchrony between the neurons and specific subsets of their afferent inputs.

## Results

Two monkeys performed a demanding shape-tracking paradigm (18,19) (Figure 1A) in which two similar visual stimuli, one attended and one non-attended were placed close together, in order to ensure a strong need for attentional stimulus selection. On top, we independently tagged both stimuli via broad-band luminance modulations. On each frame, the luminance of each shape was changed to a random value, resulting in a rapid “flickering” of the stimuli. These fast luminance fluctuations were not necessary to perform the task and were not informative in any way for the monkeys. Further, the values were changed independently for both stimuli, which made the time-varying signals unique for each stimulus (Figure 1B). To characterize the transmission of the tags of each stimulus to the local field potentials (LFPs) recorded in V4, we computed the absolute value of the complex spectral coherence (SC) between the luminance modulation of the stimuli and V4 LFP (Materials and Methods). This allowed inferring simultaneously and independently to which extent both stimuli were causally influencing neuronal activity recorded in V4. After having learned the shape-tracking task, monkeys quickly adapted to the luminance modulations and performed well on the task, despite the “flickering” of the stimuli. Excluding trials with a fixation error, monkey F performed with 79.1% correct, 9.9% early and 11.0% late responses, while monkey B performed with 91.2% correct, 7.5% early and 1.3% late responses averaged over all sessions included in the analyses (monkey F: 26.9% fixation errors, monkey B: 30.3% fixation errors). V4 LFP data was acquired in 16 sessions with 1 up to 3 electrodes per session that provided 30 recording sites (monkey F: 19 sites; monkey B: 11 sites) fulfilling the inclusion criteria (see SI Materials and Methods) which were used in the subsequent analyses.

**Figure 1.**
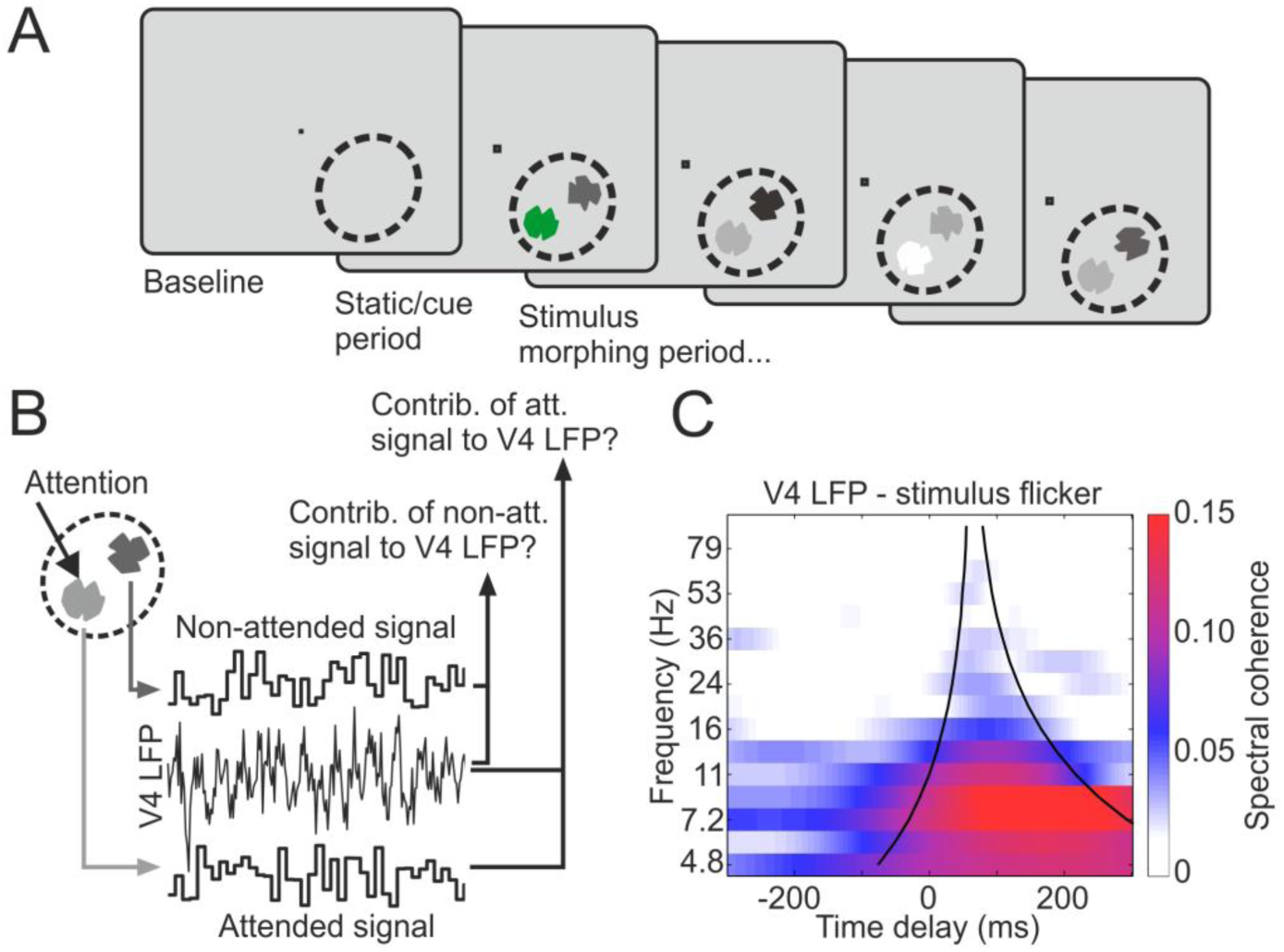
Experimental paradigm and analysis approach. (A) Outline of the task. After a baseline period, two shapes were presented together in a V4 RF (dashed lines, not visible on the monitor) and started a continuous morphing sequence into other shapes after one of them was cued (initial green coloring) to be attended. The monkeys had to respond to the reappearance of the initially cued shape. Both shapes changed luminance on each visual frame (‘flicker’), independently of each other. Note that the luminance fluctuations were much faster than the morphing speed of the shapes and that they were not relevant for the task. (B) For each trial we acquired three signals: the flicker signals of the attended and non-attended stimulus and the V4 LFP. We performed spectral coherence analysis to estimate how much each of the flicker signals contributed to the V4 LFP. (C) The average SC TdF spectrum over all recording sites from monkey B (single stimulus inside the RFs) shows the typical cone-like shape. For later site-based analyses we averaged the SC within the cone (black lines) around a time delay of 60 ms.

### Attention gates input signals to V4 LFP

The external signals, i.e. the random luminance modulations of the stimuli, contained a broad range of frequencies with their main spectral power below 50Hz (Figure S1). Time delay-frequency (TdF) representations of the SC show that specific components of these external signals causally influenced the LFP in V4, as illustrated by the average TdF spectrum of multiple recording sites (Figure 1C, single shape presentations only, all sites of monkey B included). It shows that the influence shaped the TdF spectrum in a cone-like fashion: first, it is wider for lower frequencies reflecting the temporally extended correlation structure of lower frequency signals. Second, it is shifted to the right because of the time delay of the neuronal signals. The SC between flicker signal and V4 LFP dropped with increasing frequency indicating that most of the signal components processed in V4 have frequencies below 20 Hz (Figure 1C). Note that the frequency-dependent declines in SC between flicker signals and V4 LFP are not attributable to the cutoff of the input power spectrum which reaches -3dB not before 45 Hz (Figure S1). After establishing that we could read out the external signals in V4, we used the SC measure to investigate the attention-dependent signal routing. We placed stimuli such that both were present in the RF of the local V4 population. Therefore, the neurons received synaptic input corresponding to both stimuli. These stimuli did not differ systematically with respect to their features and appeared at equivalent positions within the RF. As a result, both tagging signals were expected to contribute similarly to the V4 LFP if attention would not selectively gate the signals at the input level. An example recording site for monkey B (Figures 2A–2B, for monkey F see Figure S2), shows, however, that when attention was allocated to stimulus A, stimulus A predominantly contributed signals to the V4 LFP, whereas simultaneously very little transfer of stimulus B was observed. When attention was allocated to stimulus B, accordingly stimulus B was the major signal contributor to the V4 LFP. For each recording site, we quantified this effect by pooling the SC TdF spectra of the V4 LFP with stimulus A and B when they were attended and compared it to the pooled SC TdF spectra when the stimuli were not attended (Figures S2B and S1B): we calculated the average SC within the cone around a time delay of 60ms (see Materials and Methods). Across recording sites and monkeys, we observed strong signal gating by attention (Figure 3A), similar to the examples shown in Figure 2. From the 30 V4 recording sites displaying significant transfer of the flicker signal, 21 (9 of 11 for monkey B; 12 of 19 for monkey F) showed a significant difference between the SC of the LFP with the attended signal and the SC of the LFP with the non-attended signal. Without exception, sites with a significant attention-dependent difference in signal contribution to the LFP show that it is the attended signal that is contributing stronger to the V4 LFP. The median SC of the V4 LFP with the attended signal was more than a factor of 2 higher than that of the SC with the non-attended signal (median SC with attended signal: 0.077 range 0.048-0.152; median SC with non-attended signal: 0.036, range 0.018-0.059). For assessing the frequency-dependence of input signal gating we again computed the SC within the cone around a time delay of 60 ms for each attentional condition separately, after pooling all recording sites with a significant effect of attention. This revealed a broad-band reduction of signal transfer for the non-attended as compared to the attended stimulus (Figure 3B), with the reduction being strongest for lower frequencies, where signal transfer is strongest.

**Figure 2.**
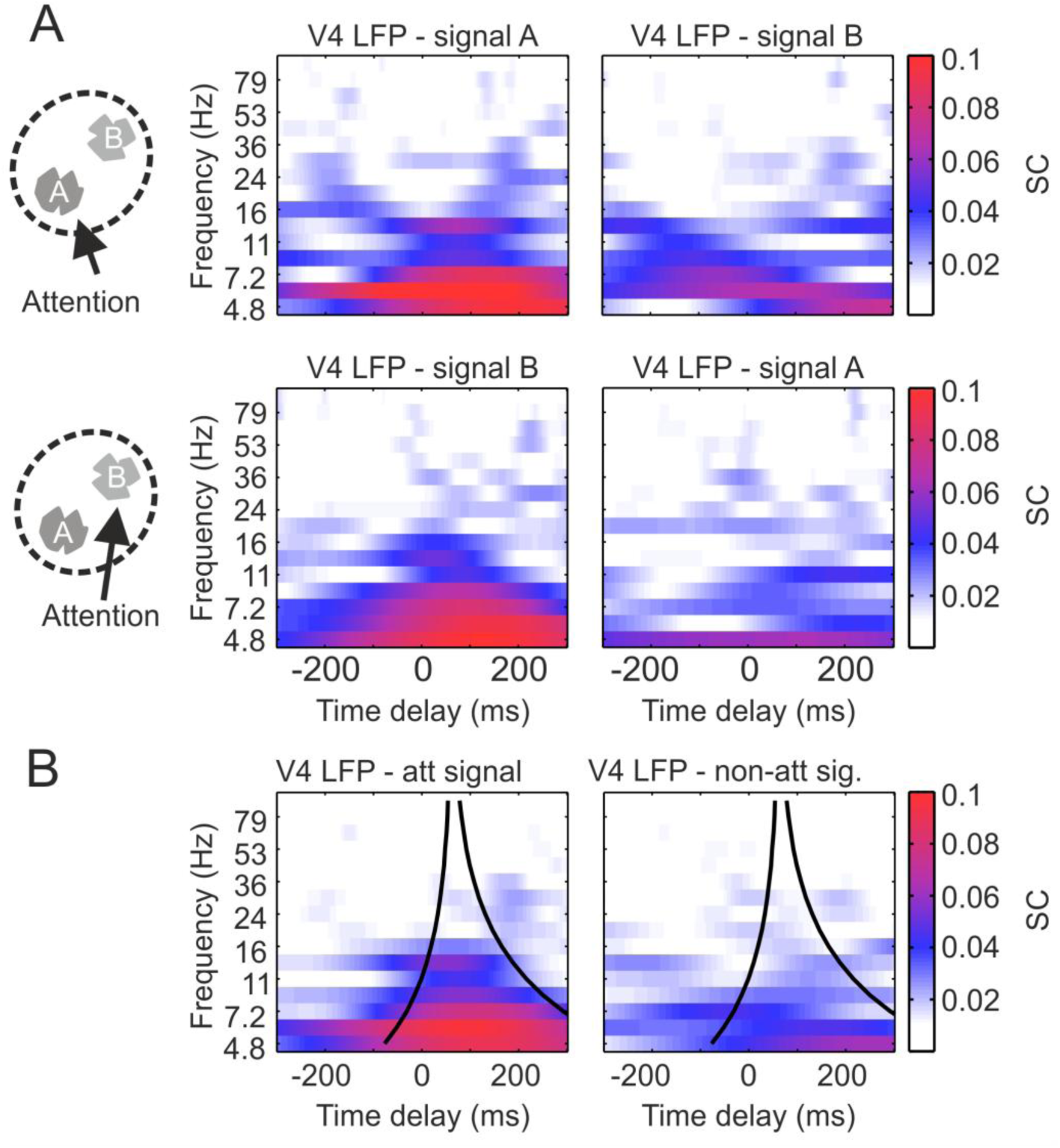
Example recording site of attention-dependent signal gating. (A) Time-delay frequency representations of spectral coherence for one exemplary recording site of monkey B. When attention was allocated to stimulus A, the SC between the V4 LFP and the flicker signal of A was high, and low between the V4 LFP and the flicker signal of B (upper row). This effect reversed when stimulus B was attended (lower row). (B) Same recording site as in (A), but data for the two attended signals (left) and the two non-attended signals (right) were pooled. Example graphs for monkey F are shown in the SI (Figure S2).

**Figure 3.**
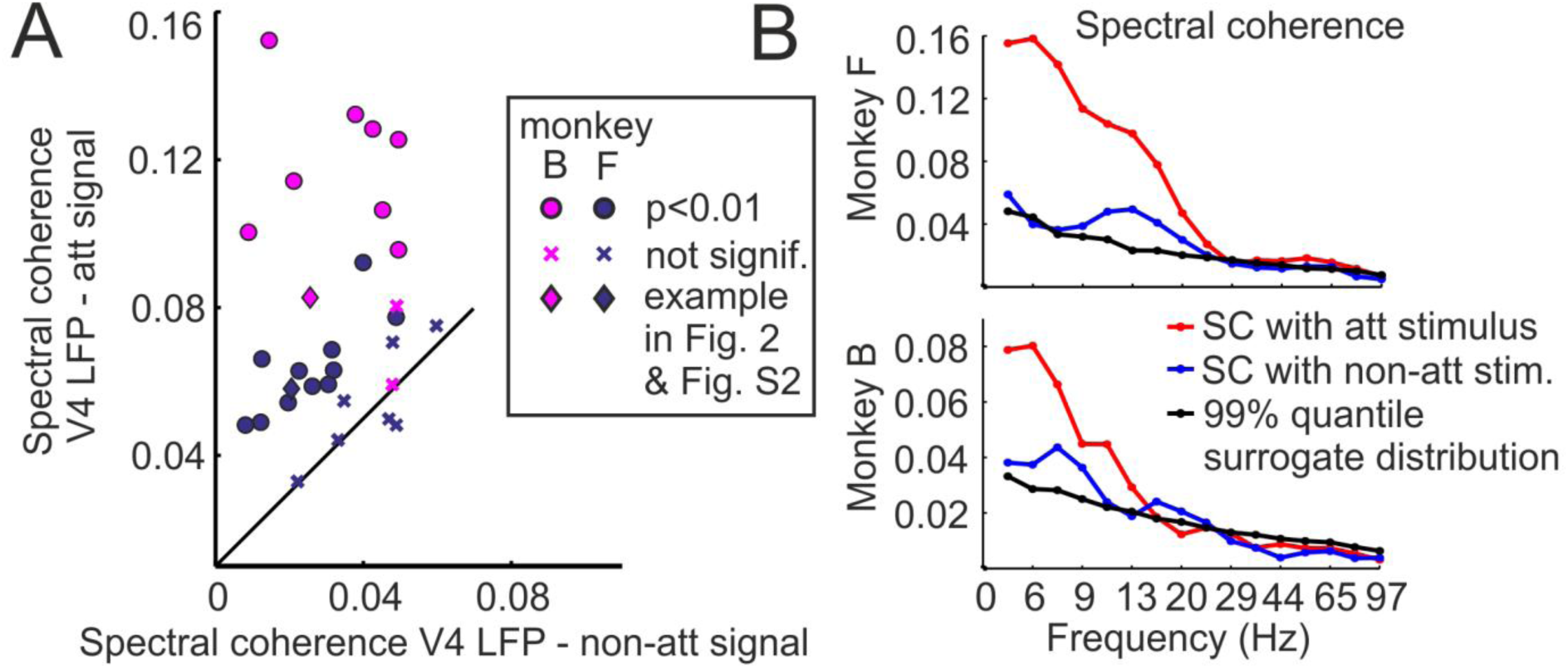
Attention dependent signal gating across recording sites and its frequency dependence. (A) Each dot, cross and diamond represents a single recording site. Dots show recordings sites with a significant (p<0.01) attentional effect, whereas crosses are not significant. Diamonds are the examples depicted in Figure 2 and S2. Magenta and blue symbols depict recordings sites of Monkey B and F, respectively. (B) The plots show the profiles of SC after pooling V4 sites with a significant gating effect (dots and diamonds in Figure 3A). The greatest differences between the SC with the attended stimulus (red lines) and the nonattended stimulus (blue lines) occur for the lowest frequencies. SC values above the black lines are significantly different from zero.

### A model explaining routing of signals

With essentially identical task requirements we recently demonstrated that selective attention goes hand in hand with highly selective gamma-band synchronization between V4 and V1 (19). In short, we showed that receiver neurons in V4 synchronize with only one of multiple input populations in V1: the population representing the attended stimulus. This suggests that selective inter-areal synchronization could underlie selective stimulus processing as observed in the present experiments. Based on these results, we built a simple model to critically test if the idea of signal routing by synchrony can explain the experimental findings (see SI Materials and Methods). Figure 4A presents the structure of this model, consisting of two input populations receiving two separate flicker signals and projecting to one common population representing a local group of neurons in V4. Each population was subjected to gamma oscillations and we simulated gamma synchrony between populations to be in or out of phase. Interestingly, we found that the SCs between the flicker signals and the simulated V4 LFP were fully consistent with the SCs calculated for the experimentally recorded V4 LFP. In the model, the “attended” signal, for which the sender population was in phase with the receiver population, was reflected much stronger in the simulated V4 LFP (compare Figure 4B with Figure 2 and Figure 3B). While the model results suggest that signal routing by synchrony can explain the observed attentional gating findings, it also predicts how the SCs between the input population’ s LFPs and the flicker time series as well as between input population’ s LFPs and the V4 LFPs would look like (Figure 5A - 5B). Further, it allowed us to test the adequacy of two different scenarios for suppressing the influence from the non-attended stimulus on V4 activity (20). The first scenario assumes a strict anti-phase relationship between gamma activity in V1 and V4, while the second scenario assumes a random phase relationship. In the model, we quantified the consequences of both possibilities on signal gating strength and the ratio of the synchrony between V4 and its two V1 input populations. In particular, we introduced a control parameter *μ* which allowed a continuous transition between the extremes of having either perfect anti-phase (*μ* = 0) or perfect random phase relationship (*μ* = 1). For example, *μ* = 0.4 would represent a situation in which in 60% of the duration of a trial, V1 is in antiphase with V4, while taking a random phase relationship in the remaining time. To analyse the relation between selective synchronization and selective transmission of the simulated stimuli, we investigated the gating strength *G* (the ratio between the SCs of the simulated V4 LFP and the attended and non-attended signals) and the synchronization ratio *S* (the ratio between the synchrony of the V4 LFP with the input populations representing the attended and non-attended stimuli) in dependence of *μ* (see SI Materials and Methods). As expected, a perfect anti-phase relation between an input population and V4 caused the strongest attenuation of the non-attended signal, leading to the highest value of *G* for *μ* = 0. Also, the synchronization ratio *S* was lower for perfect anti-phase synchronization (at *μ* = 0) than for a random phase relation between V1 and V4 gamma activity. Surprisingly, the synchronization ratio *S*(*μ*) was found to be not monotonous, but rather to exhibit a maximum at intermediate values of *μ*. This feature is caused by two counteracting effects contributing to the denominator of *S*, which is given by the SC between the V4 LFP and the input population representing the non-attended stimulus: For *μ* = 0, there is only anti-phase coherence resulting in a high SC and therefore a low *S*. For *μ* = 1, there is a mixture of on average 50% episodes with anti-phase and 50% episodes with in-phase coherence between the signals. If their associated amplitudes would be comparable, anti-phase and in-phase contributions would normally cancel each other when averaged in the SC. However, during in-phase intervals both input signals representing the attended and the non-attended stimuli are aligned and add on top of the V4 activity, hence causing the in-phase coherence contribution to be stronger than the contribution of anti-phase coherence. Taken together, with increasing *μ* the SC in the denominator of *S* first shrinks until the increasing amount of random in-phase episodes result in some recovery because of their larger V4 signal amplitudes. For our experimentally observed mean of about *G* = 2 (Figure 3A), the model predicts that the input population representing the non-attended stimulus will assume a gamma phase-relationship with V4 activity that is a mixture of being in perfect anti-phase and in a random phase (*μ* = 0.4…0.5).

**Figure 4.**
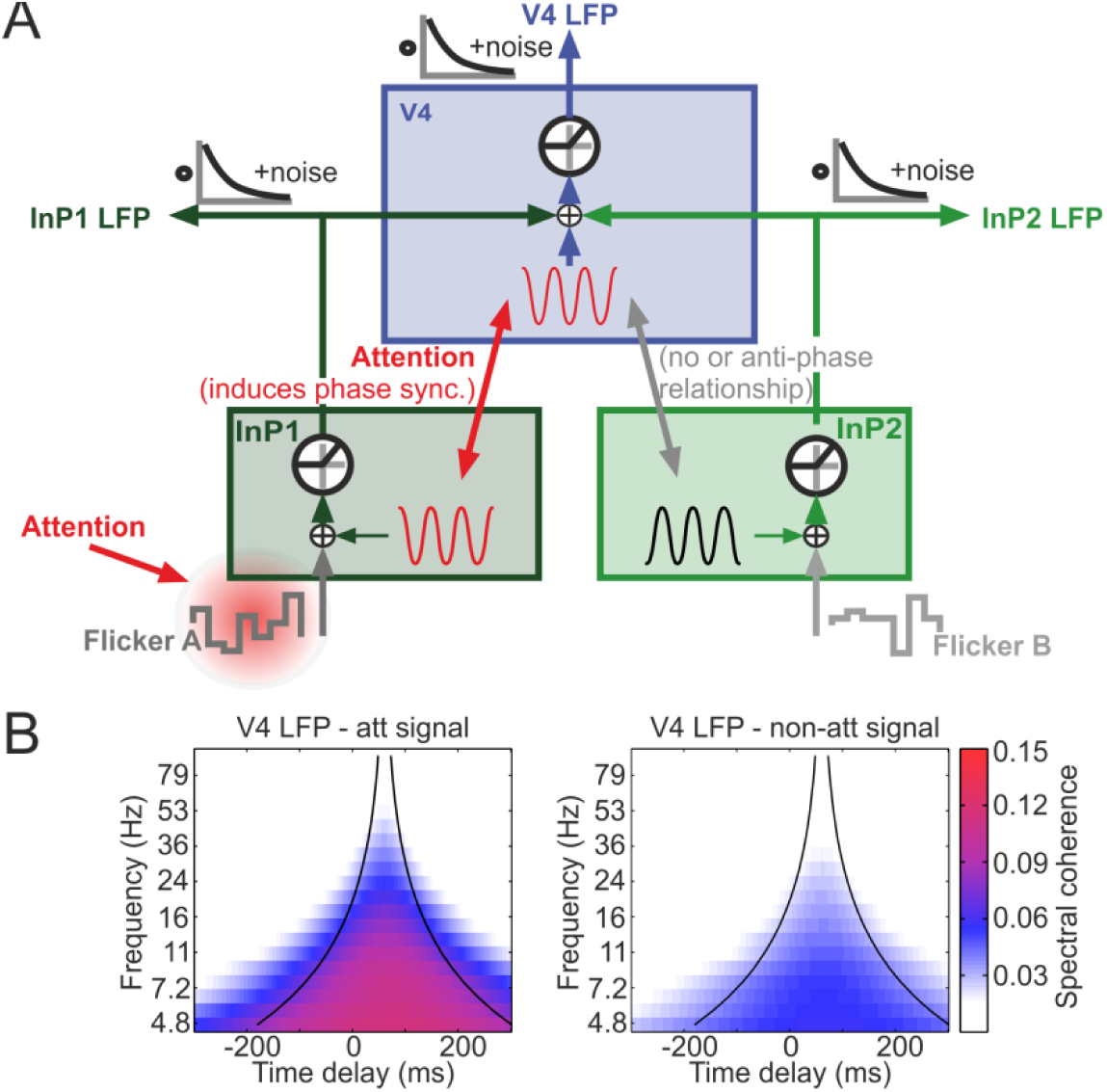
A model of signal routing by synchrony reproduces gating characteristics as observed in the experiments. (A) The model consists of three neuronal populations represented by threshold linear elements (circles). One input population such as V1 receives the flicker signal A as feedforward input, and a second input population is driven by the flicker signal B. The third population represents V4 which receives a superimposed signal from both input populations. Each population contains a noisy gamma oscillator whose signal is added to the feedforward input. LFP measurements are modeled by convolving the neural signal with an exponentially decaying kernel. Both internal dynamics and external observation are subject to additional, independent noise. Attention was modeled by synchronizing the V4 gamma oscillator with the gamma oscillator of the population receiving input from the attended stimulus. The gamma oscillator of the other input population, processing the non-attended stimulus, is set into anti-phase (or to a random phase) with respect to the V4 population. **(B)** The model quantitatively reproduces the SC in the attended (left) and non-attended (right) conditions (compare to experimental data in Figure 3 and S2).

**Figure 5.**
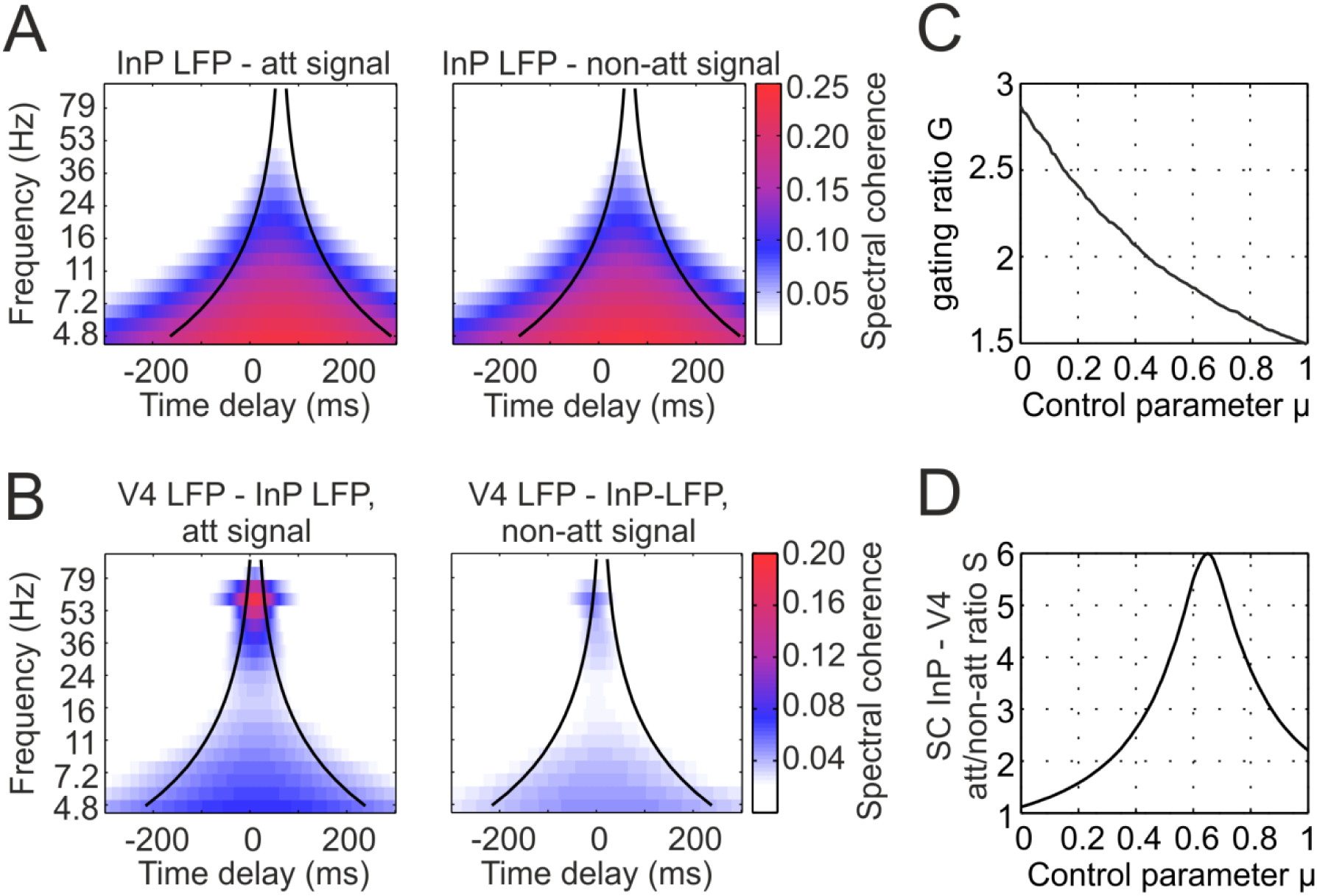
Predictions of the model. (A) The model predicts that the SCs of input populations with the flicker signal do not depend on the attention condition. The cone is centered around an assumed delay between flicker signal and input populations of 50 ms. (B) SC between input populations and simulated V4 activity is strong in the gamma range and modulated by attention. The cone is centered around an assumed delay of 10 ms between input populations and V4. (C) Predictions of the model for the gating strength (gating ratio G, left) and the attentional modulation of gamma band synchrony between input populations and V4 (synchronization ratio S, right) in dependence on the parameter *μ*. *μ* controls the randomness in the phase relationship between V4 and the input population representing the non-attended flicker signal.

For this regime, we would predict a synchronization ratio S between 2.5 and 3.8, which is well in line with the experimental observation of gamma phase synchronization between V1 and V4 being up to four times stronger in the attended condition (19).

## Discussion

We investigated if neuronal signals arriving in V4 are gated at the level of V4 neurons synaptic inputs or if all signals enter local processing and attention rather modifies the local computations by modulating the neuronal outputs. To this end we simultaneously quantified the availability of arbitrary, time-varying signals of multiple stimuli in V4. Signal availability in V4 was quantified by measuring the coherence of the V4 LFP with the broad-band luminance modulations which were superimposed on the stimuli. We found that the V4 LFP was predominantly coherent with the attended stimulus’ luminance fluctuations, even though both the target and the distracter stimulus were present within the V4 RF simultaneously and they did not have systematically different features. This result shows attentional gating for arbitrary signals and implies that attention-dependent, modified stimulus processing in V4 is based on a selective gating of the synaptic inputs as opposed to attentional selection occurring only after concerted entrance of the afferent signals, thus putting constraints on putative mechanisms underlying signal routing. Using the spectral coherence measure also allows determining the response function of the LFP to transient changes in stimulus brightness and hence to explore signal routing and its underlying mechanisms in more detail. We found that the V4 LFP could follow mainly flicker components below 20 Hz and showed largest coherence with the lowest signal frequencies, where also the attentional signal gating effect was strongest. A minimal model which implemented signal routing by selective gamma-band synchronization between neuronal populations (21,22) replicated both the observed characteristics of signal transfer to V4 and the attentional signal input gating effect. The model findings thus suggest that selective gamma-band synchrony may mechanistically implement the attentional gating of afferent input signals revealed by our experimental results. A vast amount of studies have shown that allocating attention to a stimulus accompanied by one or more distracters modifies the strength of neuronal responses to the stimuli (4–8,23,24)(25,26). However, unclear remained if attention is capable of switching neuronal processing between incoming signals, because the aforementioned studies could not show the fate of the afferent signals in local neuronal populations independently and simultaneously for all stimuli. Furthermore, the intra-cortical studies demonstrating attention-dependent processing required stimuli with markedly different features like different orientation or direction of motion in order to evoke clearly distinguishable mean rate responses. Instead, we used differential tagging of the neuronal signals of the two stimuli by a behaviorally irrelevant stochastic temporal structure of stimulus luminance. This in turn enabled us to use stimuli without systematic differences in their basic features. Consequently, neuronal responses to each of the stimuli were very similar. The highly selective processing of the attended stimulus’ signals in the present experiment did not depend on a difference of stimulus features between target and distracter stimuli. This constrains models set out to explain such attention-dependent selective processing. As attention-dependent modulations in upstream populations are typically very weak (6,27) and in this study unlikely (19), the observed selective stimulus processing depended predominantly on mechanisms acting in V4. Several models of selective attention implement a modulation of neuronal responses by changing the neuron’ s output gain. This may be caused by specific top-down inputs selectively affecting groups of neurons defined by their feature selectivity including their RF location (12,16). In the present study, the two stimuli resided within the same V4 neurons RF and do not drive distinct subsets of feature specific neurons in V4. Thus, in contrast to previous studies relying on stimuli with clearly dissimilar features selection of specific, anatomically or functionally defined, subsets of V4 neurons for attention-dependent modulation of their output gain was implausible. On top, when relying on output gain modulation, because of the stimulus similarity, both afferent signals would be expected to enter into local processing in V4 to a very similar extent. On the contrary, the clear predominance of signals caused by the attended stimulus in our experiment suggests that V4 neurons’ output gain modulation does not provide a straightforward explanation for the observed attention-dependent processing. An alternative mechanism is differential modulation of the inputs. This mechanism biases the effectiveness by which specific subsets of afferent synaptic input signals are transmitted to V4 neurons in favor of those signals caused by the attended stimulus. This does not imply a change of spike rates in the neuronal population delivering these afferents. Such selective input gating has been proposed by several models implementing biased competition (8–10) and may implement multiplication of input signals assumed by divisive normalization models of attention (24,28,29). Furthermore, a modulation of the efficacy of inputs to V4 neurons has been suggested to explain the lack of latency effects despite of average firing rate increases with attention (30). The neurons in such input gating models are much stronger influenced by the signals of the attended stimulus. Correspondingly, neurons in this study would be expected to preferentially express the temporal tagging patterns imposed on attended stimuli, despite of the absence of differences between target and distracter stimuli, which is well in line with the present observations. Our results therefore indicate that attention can already gate afferent neuronal input signals very effectively, as is assumed in models using selective input gating. This raises the question how such differential effective gain modulations for specific subgroups of inputs are achieved. Recent observations of selective modulations of gamma-band synchrony between V4 neurons and subgroups of their V1 input (19,31) suggest that signal routing by synchrony may serve as a corresponding mechanism. The minimal model based on this concept does not only replicate the observed attentional gating effects, but also explains the spectral transfer characteristics of the signal. Attentional allocation was solely implemented by selective gamma-band synchrony and did not require modulation in the activity of the upstream population nor a specific stimulus selectivity of the V4 population for one of the stimuli. Hence, also the model supports the hypothesis that attention can differentially gate signals already at the level of afferent synaptic input from upstream cortical areas and does not need to rely on feature differences between the stimuli, nor on output gain changes. Moreover, for the observed averaged gating ratio of about *G* = 2, the model delivers the prediction that V1 neurons representing the non-attended stimulus will assume a gamma phase-relationship with V4 activity that is a mixture of being in perfect anti-phase and in a random phase. The synchronization ratio corresponding to this regime is well in line with the experimental observation of gamma phase synchronization between V1 and V4 being up to four times stronger in the attended condition(19). Core assumptions of the model have found clear experimental support. Attention has been shown to modulate local gamma-band synchronization (18,32–38) and recent experiments have shown that a receiver neuronal population can selectively synchronize in the gamma-band with one of multiple input populations (19,31). The latter and our current findings are in line with previous proposals (21,22) and previous theoretical work (11,39,40) proposing that such switches of gamma-band synchrony may route the information selectively throughout the neuronal processing system. The tagging technique used in the present work for tracking multiple visual stimuli is related to frequency tagging used in electroencephalographic studies (41,42) and often for BCI purposes (starting with Middendorf et al., (43)). In the latter method two or more stimuli exhibit regular flicker with different constant frequencies (but see (44)). Though the principal underlying this method is similar to the presented one, “tagging” with a broad spectrum of frequencies provides an elegant way of overcoming difficulties of tagging with a periodic signal. In particular, we avoid the problem that single frequencies should be different enough to not interfere with each other or have similar harmonics, and that stimuli might be not equally difficult to attend or to perceive. In summary, with the combination of the stimulus design and analysis method we provide a novel way to investigate neuronal information processing: tagging stimuli by broadband temporal noise enables to track the information flow through cortex and to map transmission for all frequencies simultaneously. This method allows to investigate the linear components of signal gating, and to characterize the system response at different processing stages simultaneously. Taking the experimental and simulation data together, our findings support the notion that attention can gate information flow already at the level of afferent synaptic input which may explain the previously observed changes in firing rate.

## Materials and Methods

For an extended description of the techniques used, see SI Materials and Methods

### Surgical procedures, behavioral task

Details about the surgical preparation and behavioral task have been reported previously in Grothe et al. 2012. In short, two adult male rhesus monkeys (*Macaca mulatta*) were implanted under aseptic conditions with a post to fix the head and a recording chamber placed over area V4. All procedures and animal care were in accordance with the regulation for the welfare of experimental animals issued by the federal government of Germany and were approved by the local authorities. Before chamber implantation, the monkeys had been trained on a demanding shape-tracking task. The task (Figure 1A) required fixation throughout the trial within a fixation window (diameter 1–1.5 °) around a fixation point in the middle of the screen. After a baseline period, the monkeys had to covertly attend to the one of two statically presented, closely spaced stimuli that was cued (static/cue period). Both shapes started morphing into other shapes and the cued initial shape reappeared at pseudo randomly selected positions in the sequence of shapes. The monkeys were trained to respond with a lever release to the reoccurrence of the initial shape in the cued stream. A reappearance of the initial shape in the distracter stream had to be ignored.

#### Stimuli

In order to be able to track two input signals (the two stimuli) simultaneously and independently in one output signal (the recorded V4 LFP), we used filled shapes and imposed broadband luminance fluctuations on the stimuli. For this purpose, we changed the luminance of the shapes by choosing a random, integer gray pixel value with each frame update of the display. For monkey F, the values were drawn from an interval [128, 172], and for monkey B, from the full range [0, 255], corresponding to luminance fluctuations in a range of 6.9-12.5 and 0.02-38.0 Cd/m^2^, respectively. Both shape streams had their own independent flicker time series of luminance values.

#### Spectral coherence

Central to our analysis is the estimation of the contribution of each flicker signal to the V4 LFP. To do so we first calculated a time delay (*τ*)-frequency (TdF) representation of the absolute value of the complex spectral coherence between the LFP and the flicker signals. Following wavelet decomposition of the signal, it was computed on the basis of the complex wavelet coefficients *a*_*i,k*_(*t*, *f*_*0*_) of signal *i* in trial *k*, by

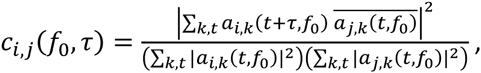

where the sums over time include all values of *t* for which *both*, *t* and *t* + *τ*, are contained within the time window of interest *T*(*k*). ā denotes the complex conjugate of *a*, and the values of *c*_*ij*_(*f*_*0*_,*τ*) lie in the range of [0,1]. Note that *τ* = 0 represents the point of zero time delay between the external flicker stimulus and its arrival in the V4 LFP. After a normalization step, we always pooled the TdF spectra of V4 LFP with stimulus A and B according to the attentional conditions. Subsequently, for a recording site-based comparison of the SC between the attended and non-attended signals we averaged the SC values in a specific TdF window. The time delay range was fixed as a frequency-dependent “cone” around an onset delay of *τ*=60ms that fits known V4 latency values (27,47) and the onset latencies observed for the evoked LFP. The cone was centered on half a wavelength after onset delay, with its borders at ±7/6 wavelength around the center to account for spread of the signal due to wavelet size (see also Fig. 1C). This wavelength dependent asymmetry of the cone was chosen based on empirical observations of a corresponding frequency dependent shift of responses in independent data taken from the stimulus onset response. The frequency range was selected for each site individually. For a given site, for both its attended and non-attended signals, we collapsed the time delay domain by averaging all SC values within the cone for each frequency bin separately. The frequency border (i.e. the highest frequency bin included) was defined as the bin preceding the first bin where both the signals were not transferred above chance level anymore (see statistical procedures).

#### Statistical procedures

For assessing significance of the spectral coherence *C* between the V4 LFP and a flicker signal, we used surrogate data. In particular, we computed a distribution of spectral coherences *C*^*S*^ between the same LFP data, but different flicker signals that were generated by using different initial seeds for the random number generator. In total, we computed 1000 surrogate samples *C*^*S*^ which were then ranked according to their value. For a (frequency-dependent) spectral coherence *C* to be significantly different from zero with an error probability below 1%, its value had to surpass the 99^th^ largest surrogate sample *C*^*S*^. For judging whether the difference *D* between two spectral coherence values *C*_1_ and *C*_*2*_ is significant, we first computed the absolute value of all differences *D*^*S*^_1,2_ between all pairs {*C*^*S*^_1_, *C*^*S*^_2_} of surrogate samples. Again, in order to be significantly different with error probability of 1%, *D* had to surpass the 99%-quantile of this distribution.

## Acknowledgments

This work was supported by the BMBF (Bernstein Group for Computational Neuroscience Bremen, grant no. 01GQ0705, Innovationswettbewerb Medizintechnik, Grant 01 EZ 0867, Bernstein Award Udo Ernst, grant no. 01GQ1106), the DFG (Priority Program 1665, grant ER 324/3), and the University of Bremen’ s Research-Focus Neurotechnology, Creative Unit I-See, and Zentrum fuer Kognitionswissenschaften. I.G. was supported by the Leibniz Graduate School for Primate Neurobiology. We are grateful to K. Thoß, R. Hakizimana, and K. Taylor for monkey care and training.

